# A CRISPR-engineered Isogenic Model Reveals Altered Neuronal Phenotypes of the 22q11.2 A-B Syndromic Deletion

**DOI:** 10.1101/2022.06.22.497212

**Authors:** Neha Paranjape, Yu-Hsiu T. Lin, Quetzal Flores-Ramirez, Vishesh Sarin, Amanda Brooke Johnson, Julia Chu, Mercedes Paredes, Arun P. Wiita

**Affiliations:** Department of Laboratory Medicine, University of California, San Francisco, CA; University of Texas Health Science Center at San Antonio, TX; Department of Neurology, University of California, San Francisco, CA; San Francisco State University, San Francisco, CA

**Keywords:** 22q11.2, microdeletion syndrome, induced pluripotent stem cell, CRISPR

## Abstract

The 22q11.2 deletion syndrome (22q11.2DS), associated with congenital and neuropsychiatric anomalies, is the most common copy number variant (CNV)-associated syndrome. Patient-derived, induced pluripotent stem cell (iPS) models have provided important insight into the mechanisms of phenotypic features of this condition. However, patient-derived iPS models may harbor underlying genetic heterogeneity that can confound analysis of pathogenic CNV effects. Furthermore, the ∼1.5 Mb “A-B” deletion at this locus is inherited at higher frequency than the more common ∼3 Mb “A-D” deletion, but remains under-studied due to lack of relevant models. To address these issues, here we leveraged a CRISPR-based strategy in Cas9-expressing iPS cells to engineer novel isogenic models of the 22q11.2 “A-B” deletion. After *in vitro* differentiation to excitatory neurons, integrated transcriptomic and cell surface proteomics identified deletion-associated alterations in surface adhesion and cell signaling. Furthermore, implantation of iPS-derived neuronal progenitor cells into the cortex of neonatal mice found accelerated neuronal maturation within a relevant microenvironment. Taken together, our results suggest pathogenic mechanisms of the 22q11.2 “A-B” deletion in driving neuronal and neurodevelopmental phenotypes, both *in vitro* and *in vivo*. We further propose that the isogenic models generated here will provide a unique resource to study this less-common variant of the 22q11.2 microdeletion syndrome.

## INTRODUCTION

22q11.2 deletion syndrome (22q11.2DS) is the most common known microdeletion syndrome, occurring in ∼1 in 1000 pregnancies and ∼1 in 4000 live births (Burnside, 2015; McDonald-McGinn et al., 2015; Panamonta et al., 2016). 22q11.2DS is characterized by range of clinical symptoms such as craniofacial abnormalities, congenital heart defects, and other developmental, cognitive and psychiatric anomalies. 22q11.2 microdeletion is a major risk factor in schizophrenia (SCZ), intellectual disability, and autism spectrum disorder (ASD) (Monks et al., 2014; Vorstman et al., 2006). The disease phenotypes are, however, remarkably variable in presentation and penetrance. While several “critical genes” at this locus have been proposed to be related to specific phenotypes (Lopez-Rivera et al., 2017; Merscher et al., 2001), no single gene is sufficient to explain the constellation of phenotypes and variable penetrance observed in 22q11.2DS.

Genomic deletions in the chromosome 22q11.2 region are caused by non-allelic homologous recombination between several repetitive sequences known as low copy repeats (LCR). The most common deletions within the 22q locus are approximately 3 Mb and 1.5 Mb in size and extend from LCR-A to D and LCR-A to B, respectively. The A-D deletion is found in ∼85% of patients and usually arises *de novo*, while the A-B deletion is found in ∼5-10% of patients and is inherited in approximately half of these cases (McDonald-McGinn et al., 1993; Morrow et al., 2018). Related to this increased frequency of inheritance, neurodevelopmental phenotypes identified in the A-B deletion may be milder than those found in the A-D deletion (Hiroi et al., 2013). However, the reason for this potential difference in patient phenotype remains unclear.

To delineate the pathogenic mechanisms of the 22q11.2 deletion, several murine genetic models have been established (Drew et al., 2011; Meechan et al., 2009; Mukai et al., 2015; Sun et al., 2018). However, while these models can recapitulate some features of the human condition, the murine chromosome 16 region does not feature perfect synteny with human chromosome 22 (Guna et al., 2015). Furthermore, particularly for neurodevelopmental effects, murine phenotypes may not reflect those in humans (Zhao and Bhattacharyya, 2018). In parallel, human-centered disease models, using induced pluripotent stem (iPS) cells derived from patients, have now come into widespread use in neurodevelopmental and neuropsychiatric research (Whiteley et al., 2022). Prior studies have indeed used iPS-differentiated neurons and neuronal progenitor cells (NPCs) (Li et al., 2019; Shimizu et al., 2022; Toyoshima et al., 2016), and, most recently, cerebral cortical organoids (Khan et al., 2020), derived from 22q11.2DS patients, to study mechanisms of this condition. However, these models are also limited due to potential genetic heterogeneity between patients, which may confound analysis of the specific impacts of the 22q11.2 deletion. In addition, essentially all described iPS models of the 22q11.2DS are derived from patients with the A-D deletion. The less common, but more frequently inherited, A-B deletion therefore remains largely under-studied.

Here we sought to address this gap in available models of the 22q11.2 deletion syndrome. We used a recently described CRISPR-based strategy (Tai et al., 2016) to generate isogenic iPS deletion clones harboring the 22q11.2 A-B deletion. Using standard methods, we differentiated these iPS clones to neuronal progenitor cells and forebrain excitatory cortical neurons. Integrated transcriptomic and cell surface proteomics identified alterations in surface adhesion and surface signaling receptors. We further leveraged emerging approaches of xenotransplantation of iPS-derived neuronal progenitor cells into the murine brain to longitudinally explore neuronal development (Real et al., 2018) and evaluate human disease dynamics in the context of an appropriate cortical microenvironment. We found that deletion-harboring NPCs showed decreased proliferation and earlier maturation to cortical neurons. Taken together, we uncover relevant neurodevelopmental phenotypes of the 22q11.2 A-B deletion while also providing a unique isogenic resource to the community for modeling this less common variant of 22q11.2DS.

## RESULTS

### Initial validation of the CRISPR-based SCORE strategy in HEK293 cells

We first sought to validate a CRISPR-Cas9-based strategy for the generation of 22q11.2 microdeletions on an isogenic background. Notably, several studies have shown that utilizing two individual sgRNA’s can lead to either deletions, duplications, or inversions at a targeted genomic locus (Kraft et al., 2015; Lupianez et al., 2015), extending up to sizes >1 Mb. However, these approaches tend to be low efficiency as they require simultaneous transduction of two sgRNA’s into the same cell, followed by DNA cleavage at two separate loci. To overcome this hurdle, for copy number variants flanked by high-homology, repetitive LCR regions, the Talkowski group described the <u>s</u>ingle-guide <u>C</u>RISPR/Cas targeting <u>o</u>f repetitive <u>e</u>lements (SCORE) strategy (Tai et al., 2016). In SCORE, only a single sgRNA targeting a homologous sequence common to both CNV-flanking LCRs is required, leading to relatively high efficiency of deletion formation via non-homologous end joining at the resulting breakpoints (**Fig. 1A**).

**Figure 1.**
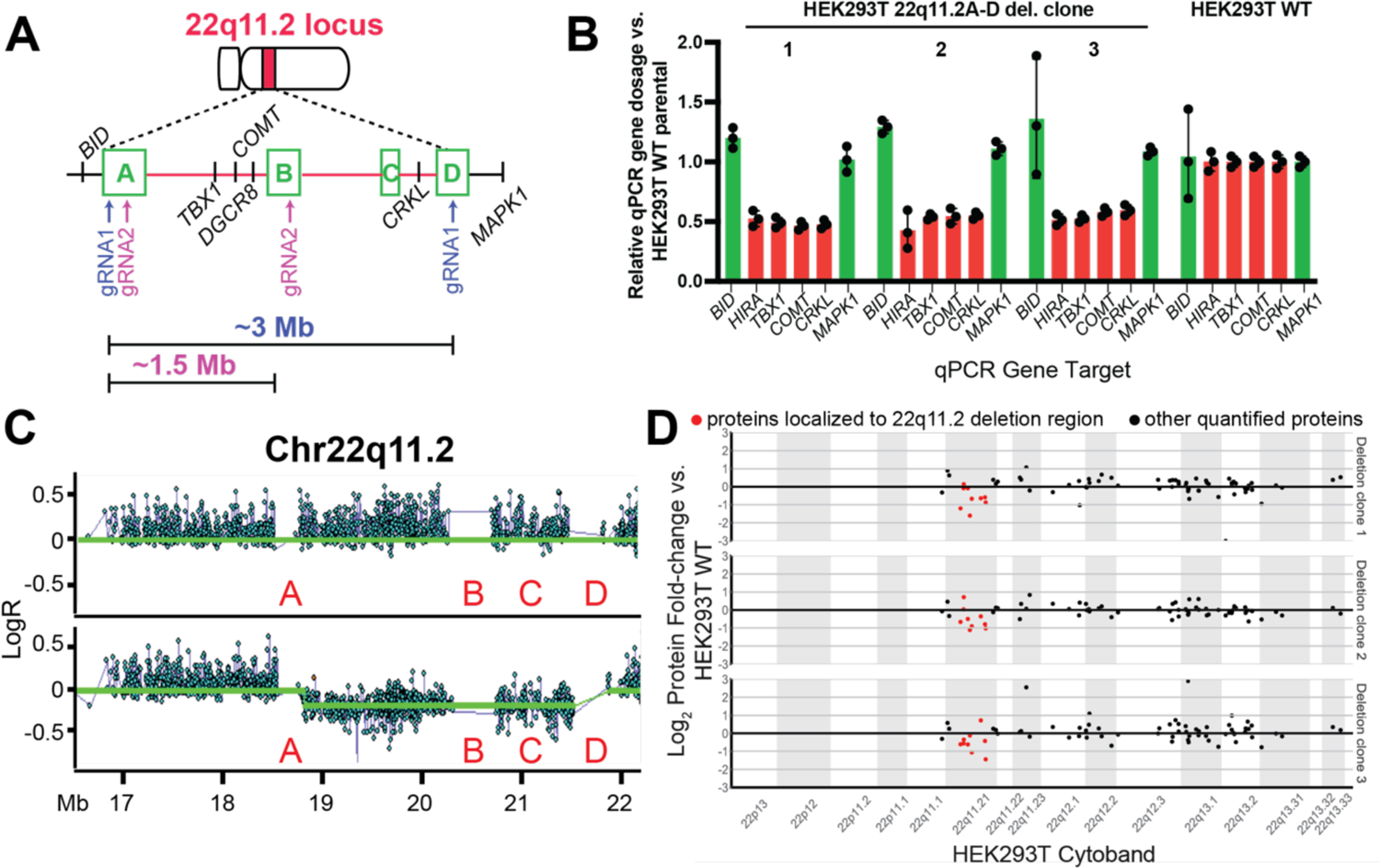
Proof-of-principle CRISPR engineering of 22q11.2 A-D deletion in HEK293T cells. **A.** Cytogenetic representation of the 22q11.2 locus on chromosome 22 in red. The 22q11.2 A-D region within the locus is depicted by dashed lines. The low copy repeats (LCRs) A, B, C and D are marked by green boxes and key genes within the region are marked. The representative breakpoints resulting from gRNA1 targeting the LCRs A and D flanking the ∼3 Mb deletion, and gRNA2 targeting LCRs A and B flanking the ∼1.5 Mb deletion, are represented by purple and pink arrows, respectively. **B.** qPCR screening of single-cell clones after genome editing with gRNA1, showing reduction in copy number of genes (*TBX1*, *COMT*, *CRKL*, *HIRA*) within the A-D deletion region in comparison to genes (*MAPK1* and *BID*) flanking the deleted region. Fold-change of genomic qPCR signal shown in comparison to parental, unmodified HEK293T cells. *n* = 3 technical replicates, +/- S.D. shown. **C.** Representative SNP microarray data from a non-edited control (top) and 22q11.2 deletion HEK293T clone (bottom) confirms deletion of the A-D region. **D.** Shotgun proteomics revealing decrease in expression of proteins encoded within 22q11.2 A-D region (in red) compared to other quantified proteins encoding on flanking regions on chromosome 22 (in black). Data shows mean-fold change per protein from *n* = 3 deletion and *n* = 3 control (non-edited) HEK293T clones, each expanded from single-cell selections.

In the original SCORE publication, isogenic iPS models of the 16p11.2 and 15p13.3 microdeletion syndromes were generated (Tai et al., 2016). Here, we sought to extend this approach to generating new models of the 22q11.2 deletion syndrome. However, it was unclear whether we could identify appropriate sgRNA’s that could successfully mediate deletions in this region. Therefore, before moving to more difficult-to-manipulate iPS models, we first aimed to validate SCORE in a highly genetically-tractable cell line model, HEK293T.

We first performed sequence analysis of the LCR “A”, “B” and “D” regions at the 22q11.2 locus (**Fig. 1A** and Methods). We used a plasmid transfection approach encoding both Cas9 and a single gRNA optimized for minimal off-target cutting using the OFFspotter strategy (Pliatsika and Rigoutsos, 2015). For these proof-of-principle experiments in HEK293T, we investigated sgRNA’s that appeared specific for the more commonly-found “A” and “D” LCRs (see Methods). After single-cell flow-sorting and colony expansion, we screened for deletions using a qPCR assay on genomic DNA for genes located within (*HIRA*, *TBX1*, *COMT*, *CRKL*) and just outside (*MAPK1*, *BID*) the deletion region (**Fig. 1B**). In initial gene dosage screening by qPCR, we achieved an encouraging success rate with 9 out of 44 (20%) isogenic HEK293T clones appearing to harbor a 22q11.2 deletion (**Fig. S1A**).

By cytogenomic SNP microarray, we confirmed that 8 of 9 screen-positive clones indeed harbored a deletion spanning the A-D LCRs at the 22q11.2 locus (**Fig. 1C**), without any evidence of additional large-scale genomic aberrations after exposure to sgRNA. Indeed, to our knowledge, this deletion (measured as ∼2.7 Mb based on unique SNP array probes; estimated size from beginning of LCR A to end of LCR D is ∼3 Mb, and commonly used as A-D deletion size (Burnside, 2015)) stands as the largest size of deletion generated using the SCORE concept.

We further investigated whether the engineered deletions carry any potential for broad cellular impacts, even in non-physiologically-relevant HEK293T cells. We specifically performed whole-cell shotgun proteomics with Label-Free Quantification, comparing 3 deletion single-cell clones versus 3 single-cell clones transduced with a non-targeting control sgRNA. Our “single-shot” analysis (see Methods) detected 9 of 43 annotated proteins mapping to the A-D CNV region. As expected, we found a decrease in the mean expression of proteins encoded in the deletion locus, compared to the mean expression of other genes on Chr22 (**Fig.1D**), as well as across the genome (**Fig. S1B**). Intriguingly, we also found alteration of similar magnitude for other proteins encoded outside the 22q11.2 region (**Fig. S1B**). While possibly indicating broader biological effects of the 22q11.2 deletion, we do note that these changes were not statistically significant based on a standard cutoff of log_2_ fold-change > |1|, and therefore may be within the experimental noise. Taken together, these results demonstrate that the single gRNA strategy can efficiently generate CNVs at the 22q11.2 locus, motivating our further studies in iPS-based models below.

### Engineering isogenic models of 22q11.2 deletion syndrome in iPSCs

After validation of the SCORE strategy at the 22q11.2 locus in HEK293T cells, we next sought to study these syndromic deletions in a more clinically-relevant background. Thus, we next aimed to model both the 22q11.2 A-D and A-B deletions in iPSCs (**Fig. 2A**). We took advantage of a well-characterized iPSC cell line from an anonymous male donor engineered to express a doxycycline inducible Cas9 protein (Mandegar et al., 2016) (CRISPRn/WTC-11). We first optimized methods for reverse transient transfection of gRNAs into iPSCs, using a co-expressed GFP marker to monitor transfection efficiency. We performed separate experiments using gRNAs we predicted using the SCORE algorithm to lead to double-stranded breaks either within the “A” and “B” LCRs, or the “A” and “D” LCRs (see Methods).

**Figure 2.**
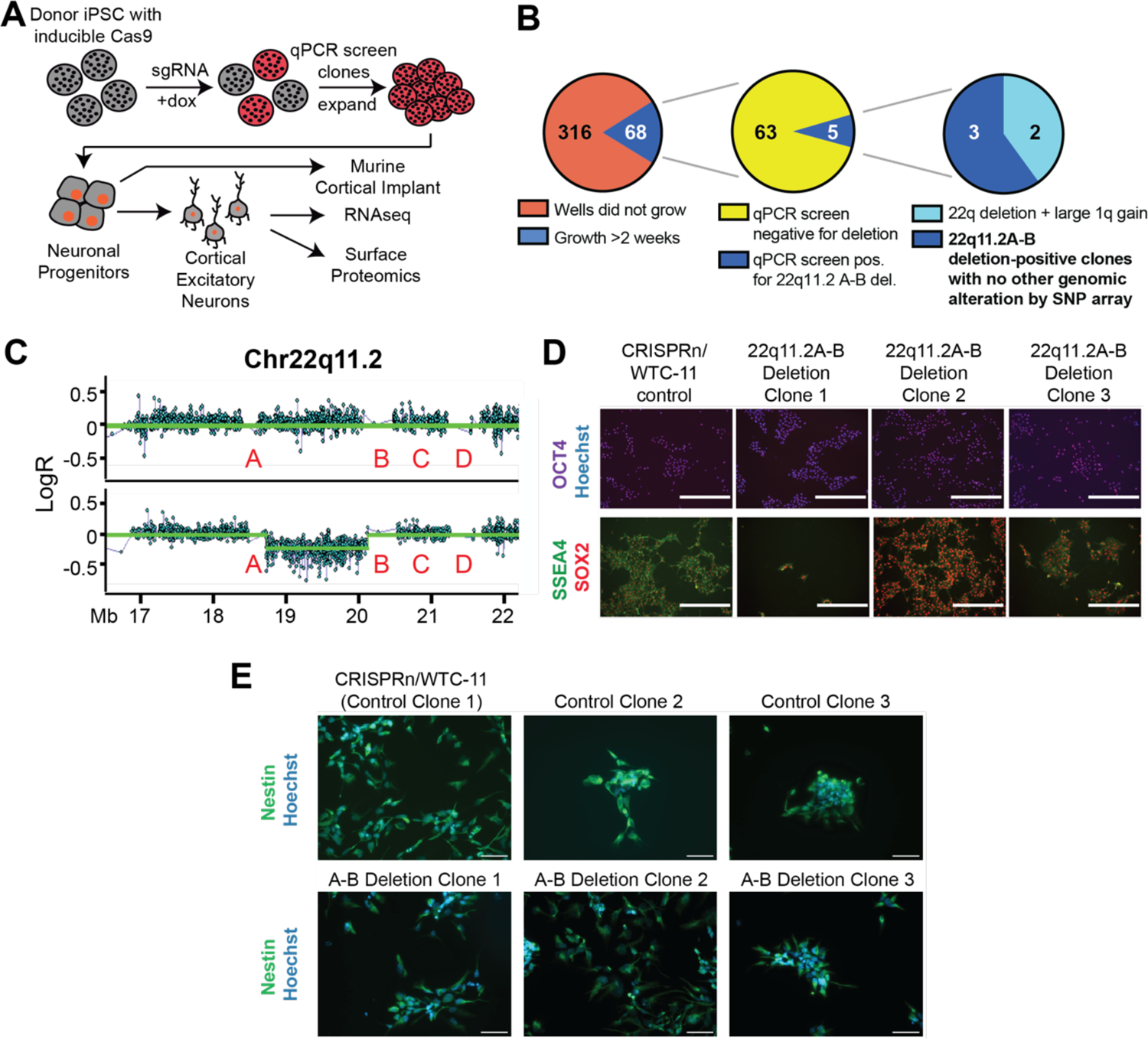
CRISPR engineering of 22q11.2A-B deletion in iPS cells. **A.** Schematic of strategy for CRISPR engineering of 22q11.2A-B deletion in iPS cells containing inducible Cas9 by using the SCORE approach, followed by differentiation into neurons and functional analyses. **B.** Single cell sorting and clonal expansion of iPS cells yielded 68 total clones expanding for >2 weeks post-sort, 5 of which screened positive by qPCR assay for the 22q11.2 A-B deletion, 3 of which were confirmed by genome-wide SNP array to harbor the 22q11.2A-B deletion and no other large genomic alterations (see **Table S1**) and were used for further study. **C.** Representative SNP microarray data from a 22q11.2A-B deletion clone (*bottom*) with comparison to control clone (*top*). **D.** Representative images of characterization of CRISPRn/WTC-11 control and 22q11.2A-B deletion iPSC clones for stem cell markers OCT4 (magenta), SSEA4 (green) and SOX2 (red). Nuclei are labeled with Hoechst (blue). Scale bar = 400 μm. **E.** Representative images of characterization of control and 22q11.2 A-B deletion NPC clones for expression of NPC marker nestin (green). Nuclei are labeled with Hoechst (blue). 40x magnification. Scale bar = 100 μm.

After single-cell sorting based on GFP+ and colony expansion, for A-B deletion clones we screened for deletions using a qPCR assay for genes located within (*TBX1*) and outside (*CRKL*) the deletion region (**Fig. S2A**). In initial gene dosage screening by qPCR, we achieved an apparent success rate of 5 out of 68 (∼8%) isogenic iPS clones potentially harboring the ∼1.5 Mb A-B deletion (**Fig. 2B**). By genome-wide SNP microarray, we further validated that 3 of these clones harbored a 22q11.2 A-B deletion and no other significant chromosomal structural aberration (**Fig. 2C** and **Table S1**).

For the designed A-D deletion, using the qPCR screening approach described in Fig. 1, we were only able to successfully generate one clone for the larger ∼3 Mb deletion, out of 35 that initially survived single cell sorting (**Fig. S1C-D**). Unfortunately, we found that even this one clone expanded very slowly, with low viability, and underwent spontaneous differentiation from the iPS stage during culture (not shown). Therefore, we were unable to use this clone for further analysis. This result suggests that iPS cells (or at least, this particular iPS line) may be less tolerant to large genomic deletions, when compared to HEK293T cells, using the same gRNA under the SCORE method. However, the ∼1.5 Mb A-B deletion appeared to be well-tolerated by the donor iPS line, with no observed deficit in iPS survival, and no spontaneous differentiation phenotype (not shown). The noted clones with this smaller deletion were thus employed for further study.

### Neuronal differentiation of SCORE-edited 22q11.2 A-B models

Given the dearth of other available models for the 22q11.2 A-B deletion, we next sought to further probe our isogenic engineered cells’ ability to reveal relevant alterations in neuronal phenotype. It is important to note that in our assays below we specifically compare our three 22q11.2 A-B deletion clones with three control iPS clones. Two of these controls are single-cell clones derived from the bulk iPS population successfully transfected with the SCORE gRNA (based on GFP positivity), but found via our qPCR screen to be negative for any 22q11 deletion (**Fig. 2B, S2A**). These isogenic control iPS clones were also verified by cytogenomic SNP microarray to not harbor any additional chromosomal aberrations (**Table S1**). We also include one clone of the parental CRISPRn/WTC-11 line, that was not transfected with an sgRNA but was selected via the same single-cell cloning process. For the former two clones, these controls have endured identical experimental perturbations to our 22q11.2 A-B deletions, controlling for impacts of sgRNA transfection and single cell cloning, but do not harbor the engineered chromosomal change. We expect the latter clone, derived from the parental CRISPRn/WTC-11 line, to reflect a useful unmanipulated comparator, as phenotypic divergence from the other two control clones may indicate impacts of sgRNA transfection rather than biology. We hypothesized that these three controls together would provide a broad-based comparator to assess the impacts of the engineered 22q11.2 deletion.

We first aimed to validate that successful generation of the 22q11.2 deletion did not disrupt the pluripotent phenotype of the iPS cells. We thus performed fluorescence microscopy for standard stem cell markers OCT4, SSEA4, and SOX2, observing qualitatively consistent expression between both 22q11.2 A-B deletion clones and control clones (**Fig. 2D**).

These results at the iPS level provided confidence to continue evaluating phenotypic alterations after differentiation to the neuronal progenitor cell (NPC) stage. We used a previously-validated approach of dual SMAD pathway inhibition to generate embryoid bodies, rosettes, and neurospheres prior to NPCs (Deshpande et al., 2017; Zhang et al., 2001) (**Fig. S2B-C**). The proportion of nestin-expressing NPCs was comparable between 22q11.2 A-B deletion and control NPCs, suggesting expected induction of neural stem cell identity (Lendahl et al., 1990) (**Fig. 2E**).

After this initial assessment at the NPC stage, we further differentiated these iPS-derived NPCs to excitatory neurons, using the approach described in (Deshpande et al., 2017; Zhang et al., 2001). We specifically focused on this cell type as our prior studies of patient iPS-derived neurons harboring a different syndromic microdeletion and reciprocal microduplication, at the 16p11.2 locus, lead to differences in cells of this lineage (Deshpande et al., 2017; Zhang et al., 2001). NPCs were plated approximately 4 weeks after beginning neural induction and then allowed to mature for approximately 3 weeks (see Methods), with validation of neuronal status based on Tuj1 positivity and appropriate morphology (**Fig. 3A**).

**Figure 3.**
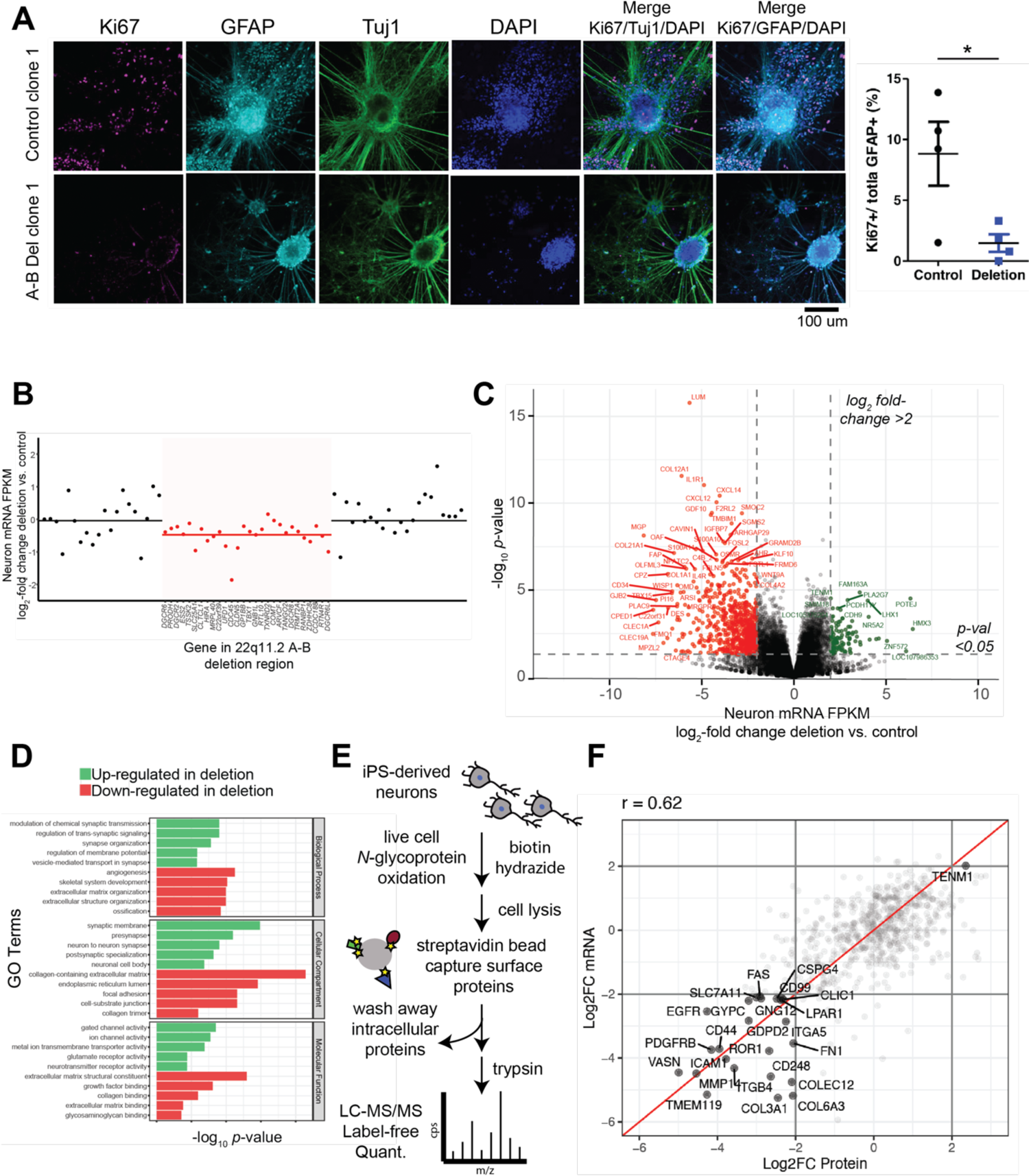
Characterizing 22q11.2 A-B isogenic deletions in iPS-derived excitatory neurons. **A:** (*Left*) Representative confocal images of control (top panel) versus 22q11.2 (bottom panel) cell lines at 3 weeks in vitro. Cells were immunostained for Ki67 (magenta), GFAP (cyan), and Tuj1 (green) expression and counterstained with DAPI. (*Right*) Quantification of the percentage of total GFAP+ cells that express Ki67 in control clones and 22q11.2 A-B deletion clone lines at three weeks in culture. Data represents *n* = 2 clones of each type (deletion clone 1 and 3, control clones 1 and 3) performed in *n* = 2 biological replicates. \**p* < 0.05 by two-tailed *t*-test. **B.** RNA-seq normalized read counts confirm that transcripts encoded within the 22q11.2 A-B deletion region (red) show lower expression in deletion clones vs. control clones when compared to transcripts encoded by flanking regions at the 22q11 locus (black). Data shows mean-fold change per transcript from *n* = 3 22q11.2 A-B deletion and *n* = 3 control clones differentiated to 3-week neurons. Solid line (red or black) indicates mean fold-change across group of transcripts. **C.** Volcano plot comparing normalized transcript abundance from RNA-seq of 3-week neurons derived from three different deletion clones compared to three different control clones. Significantly changed transcripts are labeled in red (downregulated in deletion) or green (upregulated in deletion). For significance, log_2_ fold-change cut-off = |2|; *p<*0.05 by *t*-test. **D.** Gene Ontology (GO) terms for enriched up- and down-regulated transcripts as shown in C. **E.** Schematic of surface proteomics protocol used in the study. This includes a modified (miniaturized) methodology for cell surface capture (CSC) for biotinylated proteins on ipS derived neurons that are identified by mass spectroscopy after on-bead trypsinization. **F.** Integrated surface proteomics with RNA-seq data on 3-week neurons (*n* = 3 deletion and *n* = 3 control) confirms a significant decrease in numerous surface markers in 22q11.2 A-B deletion.

Toward initial phenotypic characterization of 22q11.2 A-B deletion vs. control neurons, we quantified Ki67 in these cells as a marker of proliferation. We indeed noted a significant decrease in Ki67 in the neurons harboring the engineered deletion (**Fig. 3A**), already suggesting that the 22q11.2 A-B deletion carries some phenotypic impact in this *in vitro* system.

### Multi-omic characterization of 22q11.2 A-B deletion neurons reveals alteration of surface proteins

We next aimed to characterize the cellular-wide impacts of our engineered isogenic 22q11.2 A-B deletion in iPS-derived cortical forebrain neurons. We thus performed bulk RNA-sequencing on our neurons derived from our three 22q11.2 A-B clones versus our three controls (described above) (**Dataset S1**). As expected, we confirmed decreased abundance of transcripts encoded by genes within the 22q11.2 A-B region compared to surrounding loci (**Fig. 3B**). By DESeq2 analysis, we further found 1653 genes significantly downregulated and 830 genes significantly upregulated in the context of 22q11.2 A-B deletion (**Fig. 3C**). Gene Ontology analysis of these significantly downregulated genes demonstrated that the most prominent pathways involved cell-cell adhesion, integrin signaling, and the extracellular matrix (**Fig. 3D**). In contrast, the most upregulated biological processes and pathways related to synaptic vesicle formation and maintenance (**Fig. 3D**).

Given that many of the most downregulated genes appear to encode cell surface-localized proteins, we aimed to further perform unbiased cell surface proteomics via glycoprotein capture (Wollscheid et al., 2009) on both deletion and control neurons (**Fig. 3E**). We note that the standard version of this technique requires very high cellular inputs (>30e6 cells) (Wollscheid et al., 2009), which would not typically be feasible for *in vitro* iPS-derived neurons. However, our group recently developed an adapted “micro” approach to cell surface capture proteomics, enabling analysis of sample inputs approximately one order of magnitude lower (Ferguson et al., 2020). We therefore applied our adapted approach to ∼1-3e6 cultured neurons per condition (see Methods) and successfully quantified 637 membrane-localized proteins across all 22q11.2 A-B deletion and control samples (**Dataset S2**). We compared these results to bulk RNA-seq data and found significant concordance between quantitative surface protein expression and expression of relevant genes predicted to encode membrane-localized proteins (**Fig. 3F**) (Pearson *R* = 0.62; *p* < 0.0001). Indeed, these proteomic results confirmed that 22q11.2 A-B deletion led to downregulation of many surface localized proteins that mediate intracellular signaling including ICAM1, ITGB4, and EGFR. In contrast, we found very few significantly upregulated surface protein changes. This observation is also in accord with our RNA-seq data, where predicted significant upregulated changes are primarily expected in intracellular (i.e. non-surface) proteins, which would not be detected by this surface glycoprotein capture assay.

Taken together, these multi-omic findings suggest that the engineered 22q11.2 A-B deletion impacts membrane protein expression and localization, which could have downstream effects on intracellular signaling, cell adhesion, migration, and, ultimately, maturation.

### 22q11.2 A-B deletion impacts *in vivo* maturation of neuronal progenitors

We further sought to characterize functional impacts *in vivo* of the engineered 22q11.2 A-B deletion. While iPS-derived neuronal models are certainly powerful, findings from *in vitro* studies are limited in that they cannot represent important interactions with a relevant cortical microenvironment. Furthermore, the timeline of evaluation is limited by the tissue culture system. We were thus inspired by recently-described approaches, taking advantage of xenotransplantation of human NPCs (Paredes et al., 2022). Notably, other groups have specifically used human iPS-derived cells harboring a chromosomal aneuploidy to investigate relevant neuronal dynamics (Real et al., 2018).

To study the 22q11.2 A-B chromosomal microdeletion, we first *in vitro* differentiated deletion and control iPS cells to NPCs, using the methods as described above, and implanted them into neonatal pups (P0-P4) of the NOD *scid* gamma (NSG) strain. Using stereotactic approaches we injected ∼8e4 NPCs bilaterally into the forebrain cortex of each pup (**Fig. 4A**), with a goal to perform longitudinal analysis of NPC proliferation within this physiologically-relevant cortical microenvironment. We specifically focused on analyses up to 3 months post-transplant (MPT), given prior results demonstrating higher proliferation and differentiation at these times of other human NPC xenotransplants (Vogel et al., 2019).

**Figure 4.**
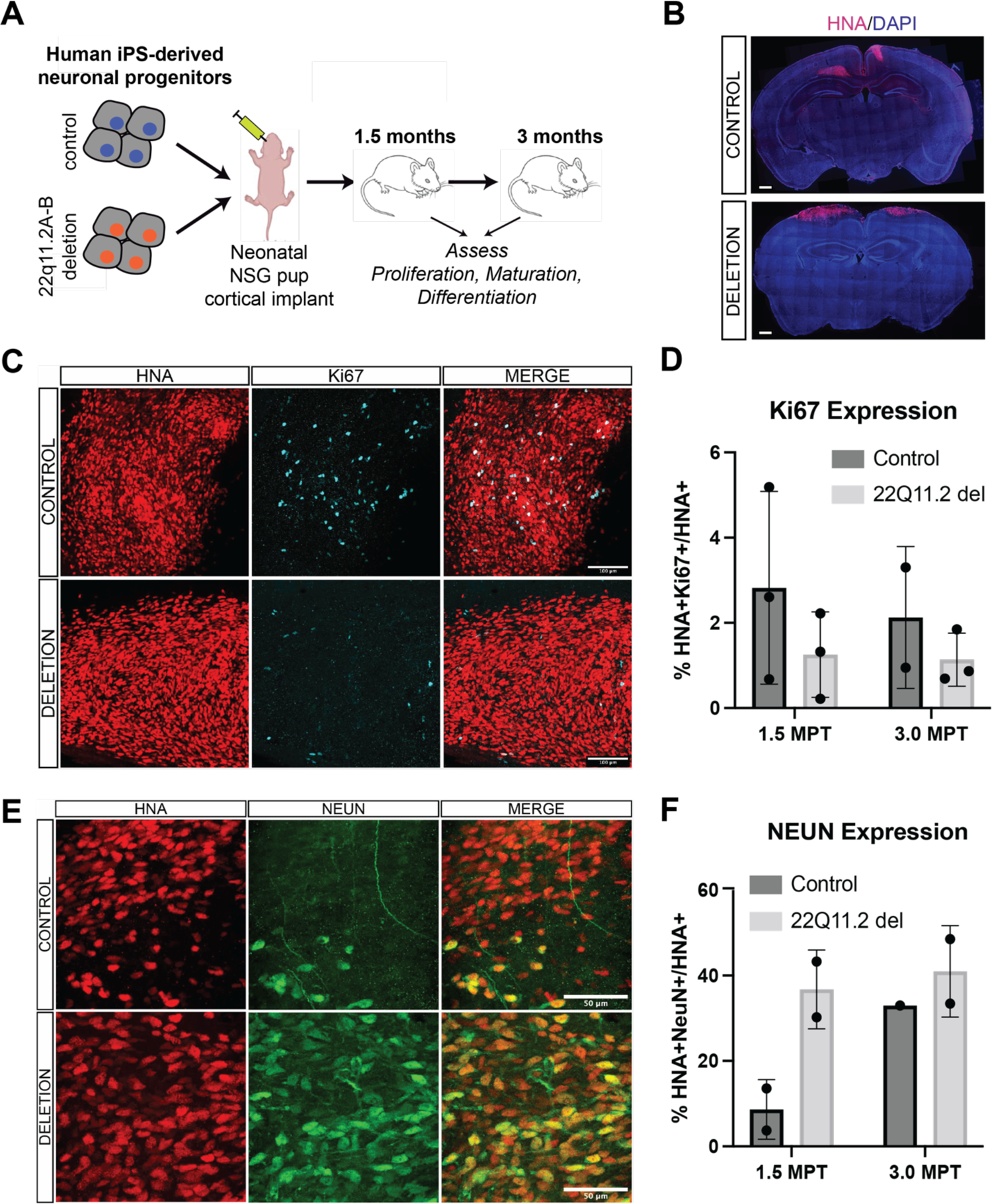
22q11.2A-B deletion causes defects in proliferation and differentiation in an in vivo xenotransplant model. **A.** Schematic of generation of NPCs for xenotransplantation injections and workflow for xenotransplantation. **B.** Coronal section indicating injection site of HNA+ cells (red) and DAPI (blue) at 1.5 MPT. Representative of *n* = 3 pups from deletion clone 3 and *n* = 4 pups from control clone 2 in this initial experiment. Scale bar = 500 μm. **C.** Representative images of transplanted HNA+ cells (red) express SOX2 (Green) and Ki67 (cyan) at 1.5 MPT in both control (top row) and 22q11.2 deletion (bottom row). Scale bar = 100 μm. Image shown from control clone 1 and deletion clone 2. **D.** Quantification of the percentage of HNA+ cells that co-express SOX2 or SOX2 and Ki67 in control (dark gray) and 22q11.2 deletion (light gray) cell lines at 1.5 MPT (left bars) and 3 MPT (right bars). Quantification performed across control clones 1-3 and deletion clones 1 and 3 (deletion clone 2 preparation had insufficient NPCs for transplant in this experiment). Each data point represents averaged data from *n* = 3 mice. *p* = ns for both comparisons by two-sided t-test. For details of quantification strategy see Methods. **E.** Representative images of transplanted HNA+ cells (red) co-express NEUN (green) in control clone 2 animals (top row) and 22q11.2 deletion clone 1 (bottom row) animals at 3 MPT. Scale bar = 50 μm. **F.** Quantification of the percentage of HNA+ cells that co-express NEUN in control (dark gray) and 22q11.2 deletion (light gray) cell lines at 1.5 MPT (left bars; control clones 1 and 2 vs. deletion clones 1 and 2) and 3 MPT (right bars; control clone 2 vs. deletion clone 1). *p*-value cannot be determined as less than three tissues are evaluated in each condition. For details of quantification strategy see Methods.

To initially validate our system, we first sought to ensure xenotransplant viability in the host mouse brain. Importantly, analysis of cortical tissue at the 1.5 MPT and 3 MPT timepoints showed viable human cells (human nuclear antigen (HNA) positivity used as a marker for human origin) at both of these time points. Notably, HNA+ cells appeared to primarily cluster near the initial injection site, illustrated both in 2-D sections (**Fig. 4B**) and Neurolucida 3-D reconstruction (**Fig. S3A**). In addition, we sought to evaluate any possible reactive or inflammatory response after xenotransplantation. Throughout neonatal NSG mouse brains we readily identified Iba1-positive macrophages (**Fig. S3B**). Notably, we found no qualitative increase in macrophage density in the regions of HNA+ cells compared to the remainder of the murine cortex; the distribution was similar in both 22q11.2DS and control NPCs (**Fig. S3B**). In addition, no visible engulfment of HNA+ cells was observed, and no ramified or activated microglia were observed in either condition. Taken together, we were encouraged that these human NPC xenotransplants appeared to readily survive over our experimental timeline and did not elicit a major murine immune response which may impact development dynamics.

We next sought to evaluate specific impacts of the 22q11.2 A-B deletion when compared to control xenotransplants. We found a trend toward decreased Ki67 of deletion vs. control clones at both the 1.5 MPT and 3 MPT timepoints, indicating that the 22q11.2 A-B deletion may cause deficiencies in cellular proliferation (**Fig. 4C-D**), and consistent with that seen *in vitro* (**Fig. 3A**). Given this result, we further probed expression of SOX2, a transcription factor known to serve as a marker of neural stem cells (Hagey and Muhr, 2014). While we did not observe any statistically significant changes in SOX2 staining, we did again observe a trend toward decreased HNA+SOX2+Ki67+ cells in the deletion vs. control implants (**Fig. S3D**). We did also observe frequent triple-positive neural rosettes in the control condition that were diminished in the deletion setting (**Fig. S3E**). Together, these findings suggest that the 22q11.2 A-B deletion may lead to decreased proliferation and more rapid loss of stem-ness within this cortical xenotransplant model.

In line with this conclusion, we next probed expression of NeuN, a marker of mature neurons. At 1.5 MPT, quantification across two deletion clones displayed 4-fold more NeuN+HNA+ cells in the transplant than controls (**Fig. 4E-F**). However, this difference appeared less pronounced at 3 MPT. Combined with the findings above, these results suggest that the 22q11.2 A-B deletion may lead to more rapid neuronal maturation of relevant progenitor cells. Together with our *in vitro* findings, these results suggest altered neurodevelopmental dynamics in the presence of the 22q11.2 A-B deletion which could affect the normal formation of the neural circuit.

## DISCUSSION

In this study we validated that the SCORE approach can be applied to generate novel, isogenic, human-centered iPS models of the 22q11.2 A-B microdeletion via CRISPR-Cas9 engineering. Profiling of these *in vitro*-derived cortical excitatory neurons, including by integrated RNA-seq and cell surface proteomics, demonstrated alterations in surface adhesion molecules and other receptors. Further functional studies using an emerging cortical xenotransplantation model demonstrated more rapid neuronal maturation than control neuronal progenitor cells *in vivo*.

We believe that our iPS-derived models will serve as a useful model system for the 22q11.2 field. Specifically, our isogenic models of the 22q11.2 A-B deletion allow for isolation of any phenotypic effects to the deletion itself, instead of the background genetic heterogeneity inherent to patient-derived models.

Prior studies have taken advantage of patient-derived iPS-based models of 22q11.2 deletions. For example, Toyoshima et al. (Toyoshima et al., 2016) and Lin et al. (Lin et al., 2016) studied iPS-differentiated neurospheres and mixed inhibitory and excitatory neurons, respectively, derived from patients with the larger 22q11.2 A-D deletion. These studies also performed transcriptional analysis of their chosen models, finding some overlap with our results, particularly in relation to changes in synaptosomal alterations and intracellular signaling pathways. More recently, Khan et al. developed iPS-derived cerebral cortical organoids from 15 patients carrying the 22q11.2 A-D deletion (Khan et al., 2020). They identified strong transcriptional signatures of alterations of excitability-related genes; we also observed changes in similar pathways (**Fig. 3D**).

Intriguingly, however, in our 22q11.2 A-B models, some of our strongest transcriptional signatures after *in vitro* excitatory neuronal differentiation related to alterations in surface adhesion molecules. This signature had not been prominently observed in the prior studies above of the 22q11.2 A-D deletion, possibly reflecting differences in 22q11.2 A-B deletion biology vs. A-D. This finding led us to complement our mRNA analysis with cell surface proteomics, identifying these alterations in cell surface molecules with an orthogonal “omics” approach.

Notably, our surface proteomic studies were inspired by prior work evaluating the surface proteomes of *drosophila* neurons, obtained via genetic engineering of a surface-tethered proximity labeling enzyme (Li et al., 2020), as well as prior work using cell surface capture on primary murine neuronal cultures (van Oostrum et al., 2020). To our knowledge, this work represents the first application of cell surface proteomics to human iPS-derived neurons. Given the critical importance of cell surface proteins in regulating neuronal migration, dynamics, and communication, we anticipate that such approaches will be highly applicable to future studies of other iPS-based neuronal disease models.

We acknowledge that a limitation of our study is that, unlike in HEK293T cells, we were unable to also successfully generate the ∼3 Mb 22q11.2A-D deletion in iPS cells via SCORE. Therefore, we could not directly compare phenotypic findings in our A-B deletion models with this more common, larger deletion on the same isogenic background. This result may reveal technical limits of the current SCORE approach for generating such large deletions in iPSCs (we note that the previous largest described was ∼2 Mb (Tai et al., 2016)). Future method optimization may enable study of even larger, clinically relevant genomic changes.

To our knowledge, our work here also represents the first xenotransplantation application of iPS-based models of 22q11.2 deletion syndrome. These longitudinal studies *in vivo*, extending over longer timescales than possible in standard *in vitro* 2-D culture, revealed a potentially relevant phenotype of more rapid neuronal maturation within the cortical microenvironment.

Notably, a prior murine model of *Tbx1* haploinsufficiency, a “critical gene” within the 22q11.2 deletion syndrome locus, also exhibited a similar phenotype of premature neuronal differentiation (Flore et al., 2017). While further study is needed, this observation in a human-derived model system may relate to neurodevelopmental phenotypes exhibited by the large majority of 22q11.2 A-B deletion carriers. In addition, further work will characterize whether alteration in any of the specific cell surface proteins we found via proteomics may play a role in the neuronal maturation phenotype, for example via altered cell-cell or cell-ECM adhesion dynamics and downstream signaling.

As our *in vivo* data demonstrates, our study was limited by relatively subtle phenotypes after murine cortex implantation, that did not typically achieve statistical significance in the quantitative expression of chosen markers. In this study, our ability to increase the number of mice evaluated was limited based on the availability of neonatal pups and technical challenges of stereotactic injections. Future studies, with larger murine cohorts, will be necessary to more fully validate functional phenotypes of the 22q11.2 A-B deletion suggested by our data.

In terms of our phenotypic characterization both *in vitro* and *in vivo*, a limitation is that we do not have any established 22q11.2 A-B deletion patient iPS lines to compare to these findings in our engineered models. However, the lack of these patient-derived specimens underscores the potential importance of the models developed here. With this isogenic cellular resource, we believe the 22q11.2 field may take finally advantage of human-centered models by which to probe this specific deletion subtype, which thus far has been largely under-studied.

## Acknowledgements

We thank Drs. Aditi Deshpande and Lauren Weiss for discussions and technical advice for completion of iPS differentiation protocols. We thank Ms. Sarah Moore for technical assistance, the Laboratory Animal Research Center (LARC) at UCSF for murine veterinary care, the Laboratory for Cell Analysis (supported by National Institutes of Health grant P30CA082103) for access to instrumentation, and Dr. Bruce Conklin for providing the CRISPRn/WTC-11 iPS line. This work was supported by a Clinical Scientist Development Award from the Doris Duke Charitable Foundation (to A.P.W.) and Roberta and Oscar Gregory Endowment in Stroke and Brain Research and DP2-NINDS 1DP2NS122550-01 (to M.P.).

## Author Contributions

N.P., M.P., and A.P.W. conceived and designed the study. N.P., Y-H.T.L, and V.S. performed cloning, differentiation and proteomics experiments. N.P., and Y-H.T.L. analyzed data. Q.R.F, A.B.J. and J.C. performed murine studies. A.B.J., J.C., and M.P. analyzed data from murine experiments. N.P., M.P., and A.P.W. wrote the manuscript with input from all authors.

## METHODS

### Cell lines

HEK293T cells were obtained from ATCC (catalog: 293T) and were cultured in standard conditions of Dulbecco’s Modified Eagle’s Medium (DMEM) (Thermo), 10% Fetal Bovine Serum (Atlanta Biologicals), supplemented with 2 mM L-glutamine (UCSF Cell Culture Facility). CRISPRn/WTC-11 cells were a kind gift of the laboratory of Bruce Conklin, Gladstone Institutes, San Francisco and have been previously characterized as having a 46,XY karyotype (Miyaoka et al., 2014). iPS lines were cultured as described below. Cell lines were routinely tested to confirm no mycoplasma contamination.

### Guide RNA design

To design the sgRNAs targeting 22q11.2 Low Copy Repeats (LCRs) with the SCORE approach, we used an in-house script that screens a contiguous DNA sequence for 20-mer sgRNA that cuts at two distinct low copy repeats within the sequence to introduce a microdeletion. We used Offspotter tool to screen out candidate guides with less than four mismatches across nontarget regions, to remove any guides with predicted off-target effects. Finally, we cross-verified that the guides target the desired regions using two different genome assemblies (GRCh38 and GRCh37) to ensure that there were no artifacts arising due to updates to the reference genome.

Sequence for 22q AB guides:

sgRNA1 sequence: GGTGCCGTCGAGAAGCGCCA

sgRNA2 sequence: GAGACGTTGAGAATGTCGCA

Sequence for 22q AD guides:

sgRNA1 sequence: GCCCTTCACTGGTTGAGTTG

sgRNA2 sequence: GTAGAAAGGGCTTTGACACG

### Derivation of 22q11.2 deletion iPSC clones

CRISPRn/WTC-11, derived from the WTC-11 line (Coriell, GM25256), is a fibroblast-derived human induced pluripotent stem cell (iPSCs) line, which expresses Cas9 under an inducible TetO promoter (Miyaoka et al., 2014). CRISPRn/WTC-11 cells were maintained on Matrigel-coated dish (Corning) with mTESR or mTESR-1 medium (StemCell Technologies) and incubated at 37 °C in a humidified atmosphere with 5% CO_2_. We used a reverse transfection method (Mirus-IT LT transfection reagent, Mirus Bio) to introduce sgRNA containing plasmid into the CRISPRn/WTC-11 cells. The gRNA plasmid also expressed EGFP. After transfection, the iPSCs were cultured on Matrigel-coated wells using mTESR medium supplemented with 10 μM ROCK inhibitor (Selleck Chemicals) for 24 – 48 hours and 2 μM doxycycline (Sigma) for 7 days. We used FACS for isolating isogenic colonies derived from single cell clones that were successfully transfected by the sgRNA plasmid. For cell sorting, the iPSCs were dissociated into a single-cell suspension with Accutase (StemCell Technologies) and resuspended in FACS buffer (D-PBS with 10 μM ROCK inhibitor). All samples were filtered through a 35-μm mesh cell strainer (Falcon) immediately before being sorted. Propidium Iodide (PI) was added to stain for viability, after gating for live (PI-) cells, the GFP+ cells (containing the sgRNA plasmid) were sorted into a Matrigel-coated 96 well plate a SONY SH800 sorter with a 100-μm nozzle under sterile conditions. A single cell was sorted into each well of a Matrigel-coated 96-well plates containing mTESR media supplemented with 10 μM ROCK inhibitor, 2 μM doxycycline and CloneR supplement (StemCell Technologies) to enhance the viability of single cells. The ROCK inhibitor was withdrawn after 24 hours and doxycycline was withdrawn after 7 days. Once multicellular colonies were clearly visible (∼7 days after sorting), they were expanded into individual wells of Matrigel-coated 24-well plates by manual picking. The iPSC colonies were further expanded into two 6 wells plates and cells were collected from one well for genomic DNA extraction and characterization for presence or absence of the 22q11.2 deletion by copy number PCR assay.

### qPCR assay to screen for 22q11.2 deletion

Genomic DNA was extracted from the iPSC colonies after single cell sorting and clonal expansion by using the Quick-DNA™ Miniprep Kit (Zymo Research 11-317A) following the manufacturer’s instructions. The DNA was diluted to 6 ng/µl for copy number screening and qPCR was performed using SsoAdvanced™ Universal SYBR® Green Supermix (Biorad, 1725272) and 1µM primer mix to amplify the target within and flanking the deleted region. The primer sequences were as follows (5’ – 3’):

*TBX1* (101 bp)

F: CCCTTACCTACCCGAGTGGA

R: AAGACGCCCATTTCTCCCAG

*HIRA* (84 bp)

F: CTGGTCACCTGATGGGCATT

R: CCCTCCCGTTCGATGATCTG

*COMT* (83 bp)

F: AGCACAGGTGGGTTTCTACG

R: AGTGAGAAAATGGAGGGCGG

*CRKL* (103 bp)

F: GTATGTTCCTCGTCCGCGAT

R: TTGGGCAGCGAGTTGATGAT

*BID* (78 bp):

F: GGCTGTGAAGGCTATGGTGT

R: AGGCTGACAGTTGAGAGCTG

*MAPK1* (70 bp)

F: CTGTGACCTCTCAGATCCTCT

R: ATCTGGCGGTTCTACAAGAGTG RPPH1 (120 bp)

F: GTGAGTTCCCAGAGAACGGG

R: TGAGTCTGTTCCAAGCTCCG

Amplification was performed by using the Step One Plus PCR system (Applied Biosystems). The following cycling conditions were used: holding stage, 98 °C for 10 min; amplification/ cycling stage, 98 °C for 15 s, 60 °C for 1 min (data collected here) (40 cycles); melt curve stage 98 °C for 15 s, 60 °C for 1 min, ramp to 98@ +0.3C/step default (data collected here) cool and hold at 40 °C for ∞. Relative copy number of genes was determined by using the ΔΔCt method. The endogenous reference gene was RPPH1 and CRISPRn/WTC was used as reference sample. For each gene, reactions were conducted in triplicate. Total reaction volume was 10µl.

### Cytogenomic SNP array

DNA was extracted from HEK293T or iPSC samples using QIAgen miniprep kit and quantified by Nanodrop (Thermo Fisher). DNA was processed for analysis by Illumina CytoSNP 850K array according to the manufacturer’s protocol. Data were processed, and CNV calls were generated with BlueFuse Multi Software v4.2 (Illumina). CNV calls by SNP array were reported using standard clinical thresholds in the University of California San Francisco Clinical Cytogenetics Laboratory, including annotation of any CNVs > 500 kb. Interpretation of clinical significance was done according to standard American College of Medical Genetics guidelines (Kearney et al., 2011).

### Derivation of NPCs from iPSCs

The control and 22q11.2 deletion carrying iPSCs were differentiated into forebrain-specific neural stem cells as previously described (Zhang et al., 2001, Deshpande et al, 2017). Briefly, iPSCs were treated with 1 U/ml dispase (StemCell Technologies) and transferred to uncoated T-25 flasks in mTESR media supplemented with 10 μM ROCK inhibitor. The cells were allowed to grow in suspension for 24-48 hours to allow formation of embryoid bodies (EBs), after which the EBs were transferred to neural media (DMEM/F-12, 1% N-2 supplement, 1% Nonessential amino acids, 2µg/ml Heparin, 1% Penicillin/Streptomycin) supplemented with small molecule inhibitors of the TGF-β and SMAD pathways, 5µM SB431542 (StemCell Technologies) and 0.25µM LDN-193189 HCl (StemCell Technologies). On day 3, EBs were transferred to Matrigel coated 6-well plates containing neural media without inhibitors for attachment and neural rosette formation. On day 11, rosettes were manually picked and transferred to uncoated culture flasks and grown as neurospheres in neural media for two additional weeks. On day 25, the neurospheres were dissociated into individual neural progenitor cells (NPCs) using Accutase and grown on PDL/Laminin coated plates for further expansion and characterization. For generation of uniform embryoid bodies, AggreWell^TM^ plates (Stemcell Technologies) were used. Briefly, Aggrewell^TM^ plates were treated with Anti-Adherence Rinsing solution (Stemcell Technologies) as per manufacturer instructions. iPSCs were enzymatically dissociated using Accutase (Stemcell Technologies). 5 – 6×10^6 iPS were seeded per single well of an AggreWell^TM^ 6-well plate. STEMdiff™ Neural Induction Medium + SMADi (Stemcell Technologies) was used per manufacturer instructions to generate NPCs from embryoid bodies (EB). EBs were cultured in Aggrewell^TM^ plates for 5 days with daily media changes, and harvested and replated on day 6 on matrix-coated tissue culture plates and cultivated for 7 days in STEMdiffTM Neural Induction Medium (Stemcell Technologies) with daily medium changes. On day 12, neural rosettes were selected using the STEMdiffTM Neural Rosette Selection Reagent (Stemcell Technologies) and replated on new coated plates. Daily media changes were continued till day 17 – 19 around which NPC outgrowth forms a monolayer between clusters. After expansion, NPCs from 22q11.2 deletion lines and control lines were immunostained for forebrain specific NPC markers such as nestin to confirm the successful derivation of NPCs from the iPSCs.

### Differentiation of NPCs into forebrain cortical neurons

The control and 22q11.2 deletion containing NPCs were plated on Poly-D-Lysine/Laminin coated plates (Corning BioCoat^TM^) for differentiation to neurons. The cells were grown in neuronal differentiation media containing Neurobasal-A, 1% Glutamax, 1% PenStrep, 2% B-27 and 1% N-2 supplements (Life Technologies) and supplements 20ng/ml BDNF, 20ng/ml GDNF (Peprotech), 200µM Ascorbic acid, 1µM cyclic-AMP and 1µg/ml Laminin (Sigma-Aldrich). The next day after plating the NPCs, a γ-secretase inhibitor Compound E (0.2µM, EMD Millipore) was added to the cells to inhibit cell division. Media changes were performed every 2-3 days. For long-term neuronal maturation (up to 6 weeks), neuronal differentiation media was switched after 2 weeks to BrainPhys™ Neuronal Medium (Stem Cell Technologies) supplemented with the same factors to promote optimal neuronal activity and maturation.

### Immunocytochemistry

The cells were grown on coated coverslips for immunostaining. The cells were fixed in 4% formaldehyde (Thermo Scientific) + 4% sucrose in PBS for 15 minutes at room temperature, washed with DPBS three times and incubated in blocking buffer (10% normal goat serum in 0.2% Triton X-100 in PBS) for 1 hour at room temperature. Primary antibodies diluted in blocking buffer was added to the cells and allowed to incubate overnight at 4°C. The next day, cells were gently washed with PBS three times and incubated with secondary antibodies for 1 hour at room temperature. The secondary antibodies were then removed, and cells were washed with PBS three times. Nuclei were stained using Hoechst 33342. The coverslips were mounted in Fluoromount-G on slides and allowed to air dry prior to imaging. Primary and secondary antibodies used are Anti-Oct4 rabbit antibody (Cell Signaling Technology, 2750, 1:400), Anti-SOX2 rabbit antibody (Millipore, AB5603, 1:500), Anti-SSEA4 mouse antibody, (Millipore MAB4304, 1:400); Anti-nestin mouse antibody (StemCell Technologies, 60091, 1:1000), Alexa Fluor 488-anti-Mouse (A11029, 1:2000), Alexa Flour 555-anti-Rabbit (A21429, 1:2000), Anti-Tuj1 (TUBB3) mouse IgG2a monoclonal antibody (Biolegend 801201, 1:1000 dilution); Anti-Ki67 rabbit polyclonal antibody (Abcam AB15580, 1:1000 dilution); Anti-GFAP chicken polyclonal antibody (Abcam ab4674, 1:1000 dilution); Goat anti-Mouse IgG2a Cross-Adsorbed Secondary Antibody, Alexa Fluor 488 (Invitrogen A21131, 1:250 dilution); Goat anti-Rabbit IgG (H+L) Highly Cross-Adsorbed Secondary Antibody, Alexa Fluor 555 (Invitrogen A21429, 1:250 dilution); Goat Anti-Chicken IgY H&L, Alexa Fluor® 647 (Abcam ab150171, 1:250 dilution). Images were obtained on EVOS FL cell imaging system, Zeiss Axioimager M1, Leica TCS SP8 Confocal Microscope, or Zeiss Spinning Disc Confocal microscope.

### Cell surface proteomics

To study the cell surface changes during neuronal maturation in the control and deletion line, the surface proteins on control and deletion 3-week neurons were labeled using a miniaturized version of the N-linked glycosylation biotin labeling method (Wollscheid et al., 2009) developed in the lab (Ferguson et al., 2020).

Briefly, 1e6 to 3e6 cells were scraped and washed twice with cold PBS, and then re-suspended in 990 μL cold D-PBS and transferred to a 1.5-mL amber tube. Next, they were oxidized using 10 μL 160 mM NaIO_4_ (Thermo 1379822) and incubated on a rotisserie at 4°C for 20 minutes. The cells were washed twice with cold D-PBS at 300x*g* for 5 minutes to remove the oxidizing reagent. For chemical labeling, cell pellets were re-suspended in 1 mL cold D-PBS followed by the addition of 1 μL aniline (Sigma-Aldrich 242284) and 10 μL biocytin hydrazide (Biotium 90060). Samples were incubated at 4°C for 60 minutes on a rotisserie followed by three more spin washes with cold D-PBS. After the final wash, supernatant was removed, and cell pellet were snap frozen and stored in −80°C until further processing for mass spectrometry. All experiments were performed in triplicates.

The labeled cell pellets were thawed on ice and lysed in 500 μL 2X RIPA buffer (Millipore 20-188) containing 1X HALT protease inhibitor (Thermo 78430) and 2 mM EDTA. Lysates were sonicated in pulses for ∼30 seconds with a probe sonicator and incubated on ice for 10 minutes. Samples were spun at 17,000g for 10 minutes at 4°C to remove cell debris. To enrich for the biotinylated surface proteins, the clarified lysates were incubated with washed Neutravidin beads (Thermo 29200) a 2-mL chromatography column at 4°C for 120 minutes.

After incubation, the beads with captured biotinylated surface proteins were washed with 5 mls of 1X RIPA + 1mM EDTA, followed by 5mls of PBS + 1M NaCl, and finally 5mls of 50mM ABC + 2M Urea buffer to remove unbound proteins. For the miniaturized cell surface capture protocol, P200 tips were packed with four C18 disks (3M 14-386-2) to create stage tips and activated with 60 μL methanol, 60 μL 80% acetonitrile (ACN)/0.1% formic acid (FA), and twice with 60 μL 0.1% trifluoroacetic acid (TFA) prior to transferring the beads to the tip using 100 μL of the 2M Urea digestion buffer. For protein digestion, 2 ug trypsin (Pierce, 90057) was added to each sample and incubated at RT for overnight digestion.

After digestion (18 – 20 hours), the pH was dropped to ∼2 with trifluoroacetic acid (TFA, Sigma, T6508-10AMP) and the peptides were allowed to bind the stage tip by gravity flow or spin filtration. The peptide mixture was desalted on the C18 stage tip by washing thrice with 0.1%TFA. Desalted peptides were eluted with 50% acetonitrile (ACN, Sigma, 34998-4L) and 0.1% TFA in LC/MS grade water and dried down completely in a speedvac. Dried peptides were resuspended in LC/MS grade water (Fisher, W64) with 2% 10 ACN and 0.1% formic acid (FA, Honeywell, 94318-250ML-F). Peptide concentration was measured using a Nanodrop (Thermo), and the peptide concentration was adjusted to 0.2ug/ul for mass spectroscopy.

### LC-MS and Data Analysis

For each replicate, 1ug of peptide was injected onto a Dionex Ultimate 3000 Nano LC instrument with a 15-cm Acclaim PEPMAP C18 (Thermo, 164534) reverse phase column. The samples were separated on a 4-hour non-linear gradient using a mixture of Buffer A (0.1% FA) and B (80% ACN/0.1% FA), from 2.4% ACN to 32% ACN. Eluted peptides were analyzed with a Thermo Q-Exactive Plus mass spectrometer.

Raw spectral data was analyzed using MaxQuant v1.5.1.2 (Tyanova et al., 2016a) to identify and quantify peptide abundance and searched against the human Swiss-Prot reviewed human proteome from Uniprot (downloaded November 26, 2018). The “match-between-runs” option was selected to increase peptide identifications. All other settings were left to the default MaxQuant values (settings as follows: enzyme specificity as trypsin with up to two missed cleavages, PSM/Protein FDR 0.01, cysteine carbidomethylation as fixed modification, methionine oxidation and N-terminal acetylation as variable modifications, minimum peptide length = 7, matching time window 0.7 min, alignment time 20 min). The MaxQuant output data was analyzed using Perseus (Tyanova et al., 2016b) and R version 3.4.0. Proteins annotated as “reverse”, “only identified by site”, and “potential contaminant” were filtered out and further filtered to remove low-quality protein quantifications. Proteins were further filtered to include only membrane-proteins or membrane-associated proteins using a manually curated list of surfaceome proteins (Nix et al., 2021). Volcano plots were generated using output from a two-sample *t*-test comparing the log_2_ transformed LFQ protein abundance values from control and deletion NPCs and neurons with a false discovery rate (FDR) set to 0.01. All proteomics results figures were produced using the R program package ggplot2.

### RNA sequencing and transcriptome analysis

RNA sequencing was done to study the RNA level changes during neuronal maturation in the control and deletion line. Control and deletion NPCs were grown in STEMdiff™ Neural Progenitor Medium (StemCell Technologies, Vancouver, Canada) on PDL/laminin coated plates prior to harvesting. Control and deletion NPCs were differentiated in neurons using Neuronal Differentiation Medium supplemented with growth factors for 3 weeks in in-vitro culture. Briefly, 1-2e^6 NPCs and 3-week neurons were harvested by scraping in an RNAse free microcentrifuge tube and washed three times with cold PBS at 300x*g* for 3 minutes. After the last wash, PBS was removed, and the cells were flash frozen in liquid nitrogen and sent to BGI Genomics (Shenzhen, China) for further processing. RNA extraction, library preparation, and sequencing were performed at BGI. RNA QC was done with Agilent 2100 Bio analyzer and Agilent RNA 6000 Nano kit. Samples were sequenced using the DNB Seq platform from BGI. For genome mapping, clean reads were mapped to reference genome (hg38) using HISAT. The average mapping ratio to the genome is 95.24% across samples. For gene expression analysis, clean reads were mapped to reference transcripts using Bowtie2 (ver 2.2.5) and expression levels calculated using RSEM (v1.2.8). The differential expression analysis was performed on counts data using DESeq2 (version 1.34.0) in R (version 4.1.2).

### Xenotransplantation

NPC CNV-containing clones and control NPCs were injected into the P0-P4 mouse cortex. NSG (NOD.Cg-*Prkdc^scid^ Il2rg^tm1Wjl^*/SzJ, Jackson Laboratories) strain mice, bred in-house at the UCSF Laboratory Animal Research Center, were used for all experiments. Injections were done bilaterally (both hemispheres) after pups were placed on ice for 4 minutes to induce hypothermia. For reproducibility, injection sites were done with precise coordinates and using anatomical landmarks such as cranial suture lines and distance from nose top. Glass needles were loaded with approximately 80,000 – 100,000 cells for each, and injection sites were found by presence of bolus cells or a needle tract scar. Mice were then warmed back up placed back into housing cage. Transcardial perfusion was done with 700 mg/kg of Avertin (2,2,2-tribromoethyl alcohol (Aldrich; T4,840-2) and tert-amyl alcohol (Aldrich; 24,048-6) by intraperitoneal injection. Perfusion was done for histological processing and only started after euthanasia induction and no response from a toe pinch from the animal. Following CO_2_ euthanasia, confirmation of death was done by visually monitoring for absence of respiration for at least one minute.

### Tissue collection

Murine brains were perfused with phosphate buffered saline solution followed by 4% paraformaldehyde and then cryoprotected in a 30% sucrose solution. Brains were embedded in OCT compound and blocks were cut into 50-micron sections on a cryostat and stored in a 0.1% sodium azide solution. Fixed brain slices were mounted to glass slides for immunohistochemistry analysis.

### Rodent perfusion and immunohistochemistry of xenotransplantation tissue

Recipient mice were sacrificed at 1.5 months or 3 months post-transplant. Mice were perfused with 4% Paraformaldehyde (PFA) solution prior to extraction of their brains and maintained in 4% PFA for 24 hours. Brains were cryoprotected with 30% sucrose to prepare for histological analysis. The brains were then cut on a Leica sliding microtome to produce coronal sections (50 μm) and kept floating in 1X PBS with .01% Sodium Azide. All histological experiments were done on floating sections. Murine tissue was washed with 1% Triton-X in TBS (washing buffer) and blocked with additional 20% donkey serum (Sigma, 9663) for 2 hours. Samples were then washed with 1% Triton-X in TBS buffer for 10 minutes and repeated 3 times. The slices then incubated with primary antibodies with denoted dilutions overnight at 4°C (anti-Iba1 goat polyclonal, Abcam ab5076, 1:200; anti-HNA mouse monoclonal, Millipore MAB1281, 1:500; anti-SOX2 goat polyclonal, Santa Cruz SC-17320, 1:200; anti-NeuN guinea pig polyclonal antibody, Millipore ABN90, 1:100). Antibodies were diluted in washing buffer with 20% donkey or goat serum (blocking buffer). Secondary antibodies (Donkey anti-goat IgG Alexa 647, Fisher A31571, 1:500; Donkey anti-goat IgG Alexa 555, Abcam ab1501350, 1:500; Donkey anti-goat IgG Alexa 594; Abcam AB150132, 1:500; Donkey anti-guinea pig IgG, Jackson Immuno Labs, 706-545-148) diluted in blocking buffer were added the following day for 2.5 hours at RT. Each experiment included a negative control to ensure no unspecific binding. Sections were mounted on microscope slides for imaging.

### Mapping of transplanted cells

50-micron sections were imaged at 10x on a Zeiss Axiovert 200M and stitched together automatically with Neurolucida software (MBF Bioscence, 2020 version). Serial sections were analyzed with 3d reconstruction with Neurolucida and transplanted cells were visualized with human nuclear antigen.

### Cell quantification

Images were acquired on a Leica TCS SP8 using a 63× (1.4 NA) objective lens. Imaging files were analyzed and quantified in Fiji software where linear adjustments to image brightness and contrast were made equivalently. Cells were counted in Z-stack images from sections stained with HNA, Ki67, NeuN, and SOX2 in 0.3 micron steps. Three to twenty representative images across a minimum of three evenly spaced and randomly sampled sections were collected for quantification at each age, and areas were based on presence of injection sites present with HNA.

## Statistics

Biological and technical replicates as performed are noted in all figure legends. No statistical methods were used to predetermine sample size. For *in vitro* and *in vivo* quantification with >3 replicates, *p*-values were obtained with two-tailed *t*-test in GraphPad Prism v9. RNA-seq volcano plot was generated with DESeq2 using standard statistical cutoffs for significance.

## Data Availability

RNA Seq data generated in the study was uploaded to GEO (Accession number GSE200225). Proteomic raw spectral data files have been deposited at the ProteomeXchange PRIDE repository (Accession number PXD032075).

---

**Table S1.** CNVs identified by SNP array in iPS clones

| iPSC line*** | Description | 22q11.2 deletion (size) | Other pathogenic* CNVs. (size) | Other VUS >500 kb** (size) |
| --- | --- | --- | --- | --- |
| CRISPRn/WTC-11 | Parental iPS line harboring dox-inducible Cas9 (control clone 1) | No | No | No |
| 22qAB_g3_B1_5 | 22q11 A-B deletion clone 1 | Yes (1.413 Mb) | No | No |
| 22qAB_g3_B1_6 | 22q11 A-B deletion clone 2 | Yes (1.413 Mb) | No | No |
| 22qAB_g3_B2-4 | 22q11 A-B deletion clone 3 | Yes (1.399 Mb) | No | No |
| 22qAB_g3_B2_1 | Control clone 2 | No | No | No |
| 22qAB_g3_B2_3 | Control clone 3 | No | No | No |
| 22qAB_g3_B1_1 | 22q11 A-B deletion, not used in further study | Yes (1.413 Mb) | 1q24.2q41 copy gain (52.454 Mb) | No |
| 22qAB_g3_B2_2 | 22q11 A-B deletion, not used in further study | Yes (1.412 Mb) | 1q24.2q41 copy gain (52.477 Mb) | No |

CNV = copy number variant
VUS = variant of uncertain significance, as defined by reporting criteria of Kearney et al. *Genet Med* **13**:680 (2011).
\*Pathogenic CNVs are defined using reporting criteria of Kearney et al. (2011).
\*\*We note that the CRISPRn/WTC-11 line (Miyaoka et al. *Nat Methods* **11**:291 (2013)) is engineered from the parental WTC-11 line, derived from a phenotypically normal male donor (karyotype 46,XY) (Coriell, GM25256). We found that CRISPRn/WTC-11 harbors two variants of uncertain significance detected by SNP array: a Yp11.2 copy loss of 2800 kb (ISCN nomenclature Yp11.2(6370460\_9170545)x0) and a 7q11.22 copy loss of 877 kb (ISCN nomenclature 7q11.2(69298131\_70158495)x1). These copy losses are present in all clones presented here. This column reflects the presence of any additional copy number changes of uncertain significance >500 kb in size, not present in the parental CRISPRn/WTC-11 line.
\*\*\*All clones except CRISPRn/WTC-11 were single-cell sorted from a bulk population that were transfected with the sgRNA designed to generate the A-B deletion and had Cas9 induced by doxycycline (see Methods). CRISPRn/WTC-11 parental iPS cells used as a control were single cell sorted from a bulk population of these cells, which were not transfected with sgRNA.

**Figure S1.**
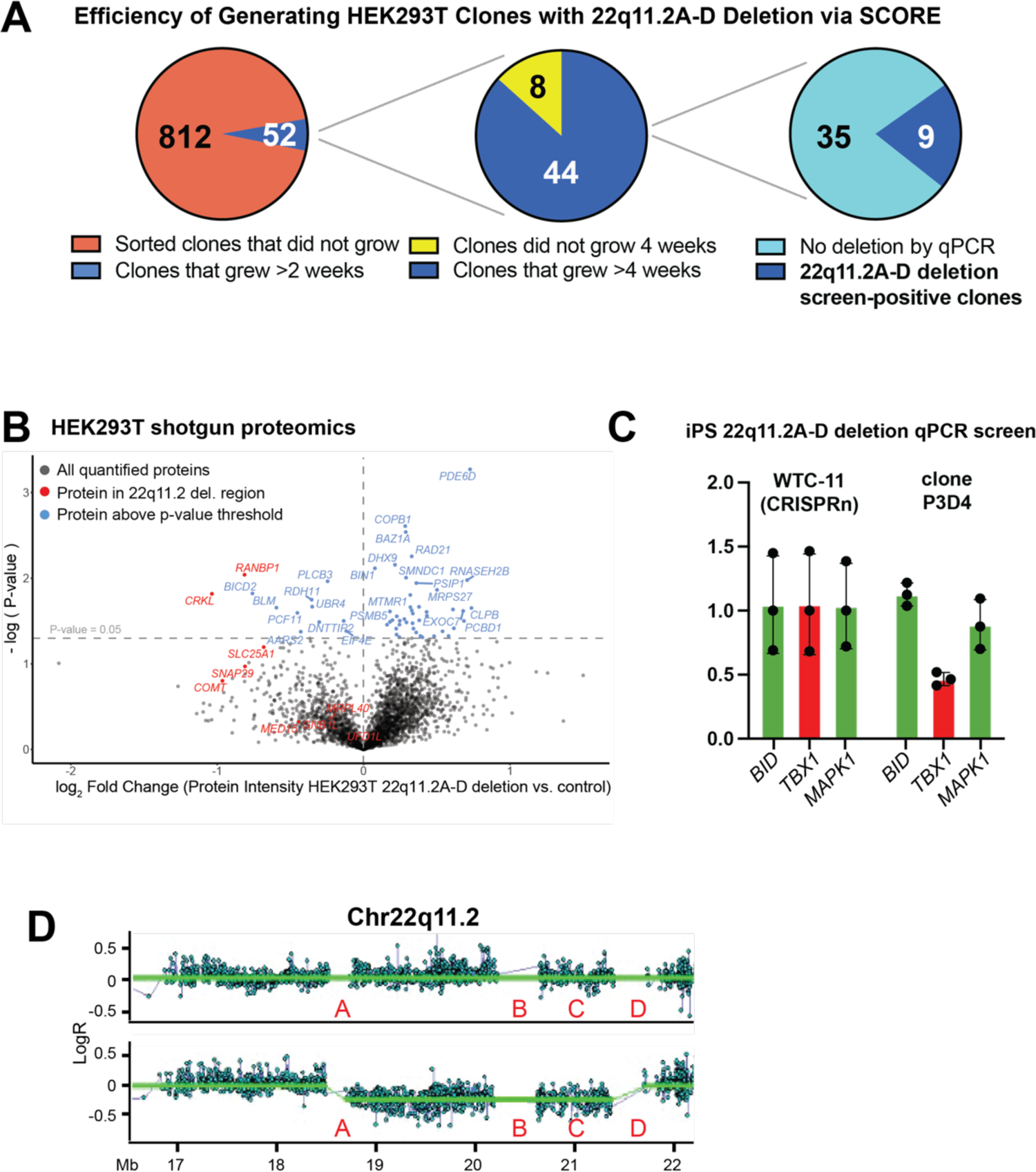
CRISPR engineering of 22q11.2 A-D deletion in HEK293T cells and iPS cells. **A.** Single cell sorting and clonal expansion of HEK cells yielded 9 deletion harboring clones from 44 total clones as determined by qPCR. **B.** Volcano plot comparing LFQ intensity of proteins in 22q11.2 deletion clones vs control clones (*n* = 3 clones each). **C.** qPCR of a single 22q11.2 A-D deletion clone (clone P3D4) showing reduction in copy number of gene *TBX1* within the deletion in comparison to genes (*BID* and *MAPK1*) flanking the deleted region. Control iPS cells (CRISPRn/WTC-11) are shown for comparison. *n* = 3 technical replicates, +/- S.D. shown. **D.** SNP microarray data of the same screen-positive deletion iPS clone confirms the presence of the 22q11.2 A-D deletion (bottom plot). CRISPRn/WTC-11 data shown for comparison (top plot).

**Figure S2.**
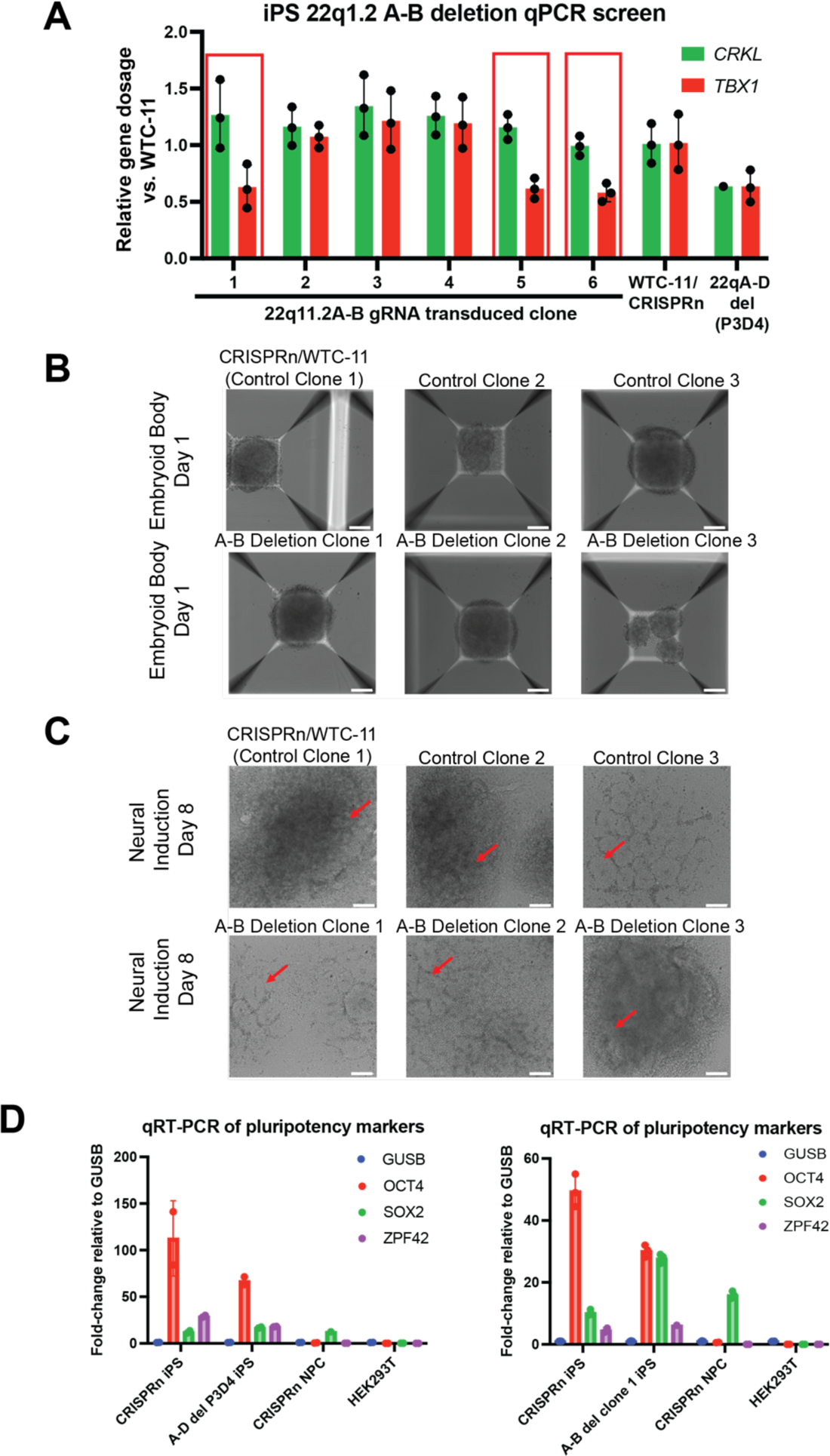
Characterizing 22q11.2 A-B deletion iPS lines. **A.** qPCR of a subset of screened 22q11.2 A-B gRNA transduced clones, showing reduction in copy number of a gene (*TBX1*) within the deletion in comparison to a gene (*CRKL*) flanking the A-B deleted region (*CRKL* located within the C-D region, see Fig. 1A). Control iPS cells and single identified clone with A-D deletion (clone P3D4) are shown for comparison. Red boxes indicate screen-positive clones (see Fig. 2B). *n* = 3 technical replicates, +/- S.D. shown. **B.** Representative microscopy images of control and deletion iPSC differentiation into embryoid bodies within Aggrewell^TM^ plates. 10x magnification. Scale bar = 100 μm. **C.** Representative microscopy images of control and deletion iPSC differentiation into neuronal progenitor cells (NPCs) via formation of neural rosettes (individual rosettes highlighted by red arrows). 10x magnification. Scale bar = 100 μm. **D.** Characterization of 22q11.2 A-D (*left*) and A-B deletion iPS (*right*) clones by qRT-PCR of standard pluripotency markers. Comparison is shown to WT CRISPRn/WTC-11 iPS and NPC clones, as well as HEK293T cells (*n* = 2-3 technical replicates per marker).

**Figure S3.**
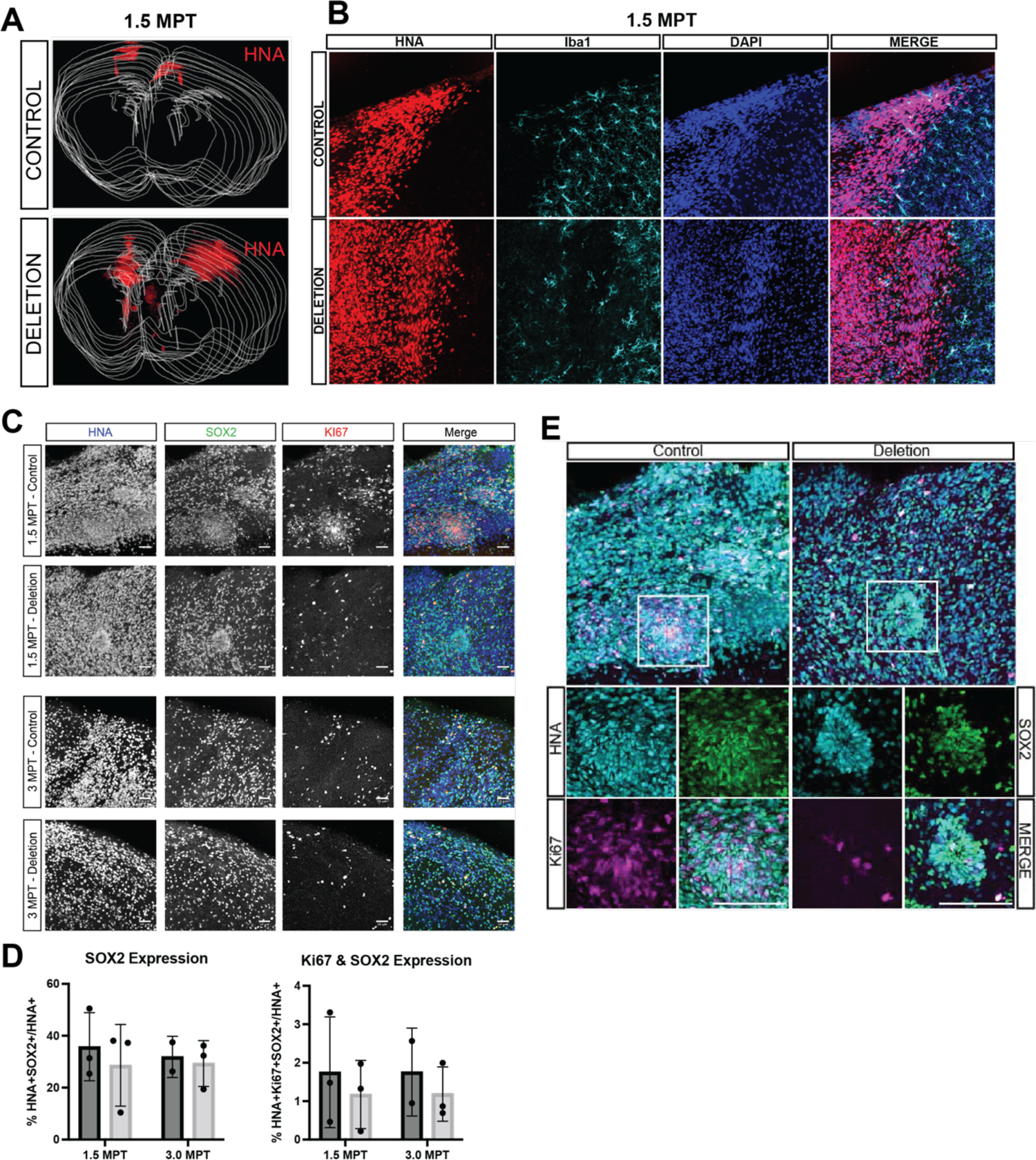
Properties of xenotransplanted control versus 22Q11.2 A-B deletion neuronal progenitor cells. **A.** Three-dimensional rendering of serial coronal sections of xenotransplanted mouse brains using control (top) or 22q11.2 deletion cell lines. Each red dot represents a xenotransplanted cell. **B.** Confocal images of control cells or 22q11.2 cell lines (identified by HNA in red) at the transplant site and surround microglia, as identified by Iba1 (cyan). Cell nuclei identified by DAPI (blue). Morphology of Iba1+ cells appear non-reactive. Representative image shown from control clone 1 and deletion clone 3. **C.** Transplanted HNA+ cells (blue) co-express both SOX2 (green) and KI67 (red) at 1.5 MPT (1st and 2nd rows) and 3 MPT (3rd and 4th rows) in control vs. 22q11.2 deletion xenografts. Representative images shown from control clone 3 and deletion clone 2. **D.** (*Left*) Quantification of HNA+ cells that co-express SOX2 in control (dark gray) and 22q11.2 deletion (light gray) at 1.5 MPT and 3 MPT. (*Right*) Quantification of HNA/KI67/SOX2 triple-positive transplanted cells in control (dark gray) and 22q11.2 deletion (light gray) at 1.5 MPT and 3 MPT. Quantification performed across control clones 1-3 and deletion clones 1 and 3 at 1.5 MPT (each data point average of n = 3 mice per clone); control clones 2 and 3 and deletion clones 1 and 2 at 3 MPT (each data point average of n = 2 mice per clone). **E.** Confocal images of injection sites containing neural rosettes formed at 1.5 MPT in control (1st row; left panel) and 22q11.2 deletion (1st row; right panel) contain transplanted cells that are triple-positive (white) for HNA+ (cyan), SOX2 (green) and KI67 (magenta). Higher magnification of neural rosettes (white boxes) show larger rosettes with more triple-positive cells present in control clone 3 (2nd and 3rd row; *left*) vs. 22q11.2 deletion clone 2 (2nd and 3rd row; *right*)

## SUPPLEMENTARY DATASET LEGENDS

**Dataset S1:** RNA-seq of three 22q11.2A-B deletion neuron clones vs. three control neuron clones

**Dataset S2:** Surface proteomics of three 22q11.2A-B deletion neuron clones vs. three control neuron clones

